# Inter-hemispheric connections modulate splitting in a computational model of the bilateral SCN

**DOI:** 10.64898/2026.04.30.722022

**Authors:** Klavdia Zemlianova, Jaden McDaniel, Alice G. Lander, Julu Nwaezeapu, Gabrielle J. Gutierrez

## Abstract

The phenomenon of splitting was originally observed in hamsters which, after prolonged exposure to constant light, exhibit two rest/wake cycles within a subjective day. Splitting is a consequence of the left and right suprachiasmatic nuclei (SCN) falling out of synchrony. While it is known that split activity is characterized by an antiphase relationship between the left and right SCN and between the core and shell within each hemisphere, the role of the commissural projections that connect the right and left SCN is not known. In the present study, we investigate the impact of the inter-hemispheric connections on the split and unsplit dynamics of a computational model of the bilateral SCN. Our model has 4 nodes corresponding to each right and left core and shell. We simulated our bilateral model under different lighting conditions and measured its period and the phase relationships among the 4 nodes. To further characterize the dynamics of the system, we performed a bifurcation analysis. We found that the bilateral model automatically splits unless entrained by bright light/dark cycles, or unless it has excitatory inter-hemispheric connections. This suggests that excitatory cross-connections may be important for freerunning behavior. We found that constant light of varying intensities transitions the model between split and unsplit activity only in very limited conditions, but the strength and polarity of the contralateral connections play a much greater role in this dynamical transition. These findings suggest that splitting may involve plasticity of the inter-hemispheric connections of the SCN.

## Introduction

### Anatomy and physiology of the SCN

The suprachiasmatic nucleus (SCN) consists of a small cluster of neurons in the mammalian hypothalamus which serves as a central pacemaker that controls and regulates the body’s circadian rhythms (Hastings et al., 2018). The SCN synchronizes internal biological clocks with environmental cues such as light and temperature, facilitating an adaptive response to the 24 hour day-night cycle (Majercak et al., 1999). The generation and maintenance of these rhythmic patterns of neural activity influence various physiological processes including, sleep-wake cycles, hormone secretion, body temperature, and feeding behavior (Dibner et al., 2010; Guenthner et al., 2014; Merrow and Harrington, 2020).

The SCN is a bi-hemispheric structure with each side containing approximately 10,000 neurons (van den Pol, 1980; Hastings et al., 2018). In each hemisphere, the SCN can be divided into two distinct regions (Card et al., 1981; Moore et al., 2002). The ventrocaudal region (core) receives direct retinal input via the retinohypothalamic tract (RHT) and does not exhibit detectable rhythms in clock gene expression and electrical activity (Moore et al., 1995; Berson et al., 2002; Antle et al., 2009; Paul et al., 2009). In contrast, the dorsal region (shell), is populated with cells that exhibit strong circadian rhythms in both clock genes and clock-controlled genes, despite the lack of direct retinal input (Colwell, 2011). Individual SCN neurons are capable of generating circadian rhythms in electrical activity as seen in vivo, in SCN tissue cultures, and in brain slice preparations (Meijer et al., 1998; Yan et al., 2005; Maywood et al., 2006; Nakamura et al., 2008; Webb et al., 2009; Bano-Otalora et al., 2021). These rhythms can be sustained in darkness and can even be recovered by transplantation of a grafted SCN into a host (Silver et al., 1996).

The connectivity within the SCN and among its constituent regions is responsible for synchronized circadian activity (Antle and Silver, 2005; Maywood et al., 2006). SCN neurons are highly interconnected through both synaptic and peptidergic signaling (Colwell, 2011; Hastings et al., 2018). GABA is released by nearly all SCN neurons and mediates inhibition as well as excitation depending on the postsynaptic response (Farajnia et al., 2014; Albers et al., 2017; Hastings et al., 2018). The core contains neurons that express vasoactive intestinal peptide (VIP) and neurons that express gastrin-releasing peptide (GRP), while the shell contains vasopressin (AVP)-expressing neurons (Antle and Silver, 2005; Colwell, 2011; Hastings et al., 2018). The core sends projections to the shell, and although there is little evidence for shell to core projections in rodent SCN, commissural fibers connecting the left and right cores and the left and right shells have been observed (van den Pol, 1980; Card et al., 1981; Pickard, 1982; Leak et al., 1999; Abrahamson and Moore, 2001; Moore et al., 2002). The commissural connections are thought to be mediated by glutamate in the mouse (Michel et al., 2013). While these commissural projections connecting the left and right hemispheres of the SCN have been observed in multiple studies, the function that such inter-hemisphere connections serve is not well established.

### Split rhythms

The clock system exhibits remarkable flexibility in response to specific environment conditions. The golden hamster, *M*. auratus, is a nocturnal creature with bouts of rest/wake activity that follow a circadian period of approximately 24 hours. Under conditions of constant darkness, the hamster’s circadian rhythms free-run as they maintain behavioral rhythmicity (Rusak, 1977). However, when exposed to constant light for prolonged periods of several days to several weeks, hamsters display a unique phenomenon called “splitting” (Pittendrigh, 1960a; Pittendrigh and Daan, 1976; Pickard and Turek, 1982). Splitting refers to the divergence of an organism’s activity patterns, creating two distinct components within a 24-hour period (Welsh et al., 2010). Effectively, the split animal experiences transitions from rest to wakefulness at about twice the frequency as in free-running. Splitting can occur in dim light, but the likelihood of transitioning to splitting is dependent on the intensity of the constant light, with brighter constant light increasing the likelihood of splitting (Pickard et al., 1993). The period of circadian rhythms lengthens as light intensity increases, and animals with longer free-running periods are more likely to exhibit splitting (Pickard et al., 1993). The amount of wheel-running activity influences the latency to split where more active hamsters have a shorter latency for transitioning to the split rhythm (Butler et al., 2012).

The critical role of the SCN in circadian regulation is demonstrated by studies where lesions to the SCN of hamsters resulted in abolishment of the splitting behavior. Unilateral damage to the SCN in split hamsters restores the unsplit rhythm (Pickard and Turek, 1982). Bilateral and unilateral lesions to areas surrounding the SCN can also abolish the split rhythm, demonstrating that split behavior is sensitive to the interactions within a complex system that extends beyond the SCN (Harrington et al., 1990).

The gradual divergence of the 24-hour rest/wake rhythm into a split rhythm led researchers to hypothesize that there are two components to the clock that may synchronize either in unison or alternately (Pittendrigh, 1960a; Pittendrigh and Daan, 1976; Rusak, 1977; Evans and Schwartz, 2024). Indeed, the extinction of the split rhythm after unilateral lesions to the SCN lent support to that idea (Pickard and Turek, 1982), and later it was observed that split hamsters have asymmetric expression of period genes between the left and right SCN (de la Iglesia et al., 2000). *Per1* and *Bmal1* are normally expressed symmetrically across the left and right SCN while alternating during a 24-hour period. In the split hamster, *Per1* expression was observed unilaterally while *Bmal1* expression was simultaneously found in the contralateral SCN, indicating an antiphase relationship between the left and right SCNs of split animals (de la Iglesia et al., 2000). The physiology of the split SCN was further refined by studies showing an antiphase relationship for *Per1* expression in the core and shell (Yan et al., 2005). c-FOS expression is also in antiphase between the core and shell of the split animal, demonstrating a correlation between electrical activity and clock gene expression (Tavakoli-Nezhad and Schwartz, 2005; Yan et al., 2005; Butler et al., 2012). Each SCN hemisphere of the split animal individually exhibits a 24-hour rhythm but each hemisphere is in antiphase with the other, leading to the 12-hour rest/wake cycles that characterize the split behavior. Extra-SCN regions, on the other hand, exhibit in-phase 12-hour rhythms in correspondence with the behavior (Butler et al., 2012).

### Modeling the SCN and split rhythms

The phenomenon of splitting in hamsters has prompted researchers to explore the existence of two distinct oscillators governing circadian rhythms, rather than a single oscillator, in mathematical models of the SCN (Daan and Berde, 1978; Kawato and Suzuki, 1980). While several variations of mathematical and computational models of splitting exist, examining the physiology of the split SCN requires a model that captures the antiphase relationship between the left and right hemispheres while also embodying the antiphase activity between the core and shell within a hemisphere. In the present study, we create a computational model that can simulate the dually antiphase activity observed in recent studies (Yan et al., 2005; Butler et al., 2012), which we term the 4-node split.

Despite the growing body of literature, significant knowledge gaps remain regarding the precise synaptic and circuit-level dynamics that facilitate splitting. To address these gaps, the present study constructs a model of the SCN from a pair of coupled gated oscillators based on the model by Carpenter and Grossberg, 1984. Our model differs from previous models by explicitly incorporating the lateralized structure of the left and right hemispheres of the SCN, allowing the simulation of the 4-node split activity and to identify critical thresholds in coupling strength that transition the model between split and unsplit activity. The results extend beyond newly hypothesized circuit mechanisms underlying split behavior; they offer insight into the broader dynamics of circadian regulation in response to environmental changes.

## Methods

### Bilateral SCN model

Our model is an extension of a classic gated pacemaker model (Carpenter and Grossberg, 1984). The original model has two nodes which are mutually coupled, forming a half-center oscillator. Each node features a gating mechanism capturing the dynamics of a neural system where the depletion of neurotransmitters reduces the effective signal transmission between nodes in the circuit. When the original model was conceived, researchers had found that there are parts of the SCN that receive retinothalamic inputs and that different parts of the SCN have distinct chemoarchitectures; however, the core and shell organization of the SCN had not yet been fully established (Moore et al., 2002). In alignment with contemporary terminology, we denote Carpenter and Grossberg’s abstract chemoregional variables as core and shell. Light is received by the *υ*_*core*_ node, simulating the retino-recipient nature of the core of the SCN. The output of the *υ*_*shell*_ node is a proxy for the animal’s activity in this model. A Fatigue signal, *F*, integrates the shell activity and feeds it back to the core.

Our model innovates upon Carpenter and Grossberg’s original model by incorporating bilateral SCN nodes (Fig. 1A). Expanding the model from unilateral core and shell nodes to bilateral core and shell nodes requires four dynamical variables (*υ*^*R*^_*shell*_, *υ*^*L*^_*shell*_, *υ*^*R*^_*core*_, *υ*^*L*^_*core*_) and their corresponding gating variables (*z*^*R*^_*shell*_, *z*^*L*^_*shell*_, *z*^*R*^_*core*_, and *z*^*L*^_*core*_). The dynamical variable for Fatigue, *F*, integrates the mean left and right shell activity. Incorporation of distinct bilateral SCN populations enabled our investigation of splitting and the dynamics that govern the transitions between split and unsplit activity in the SCN.

**Figure 1.**
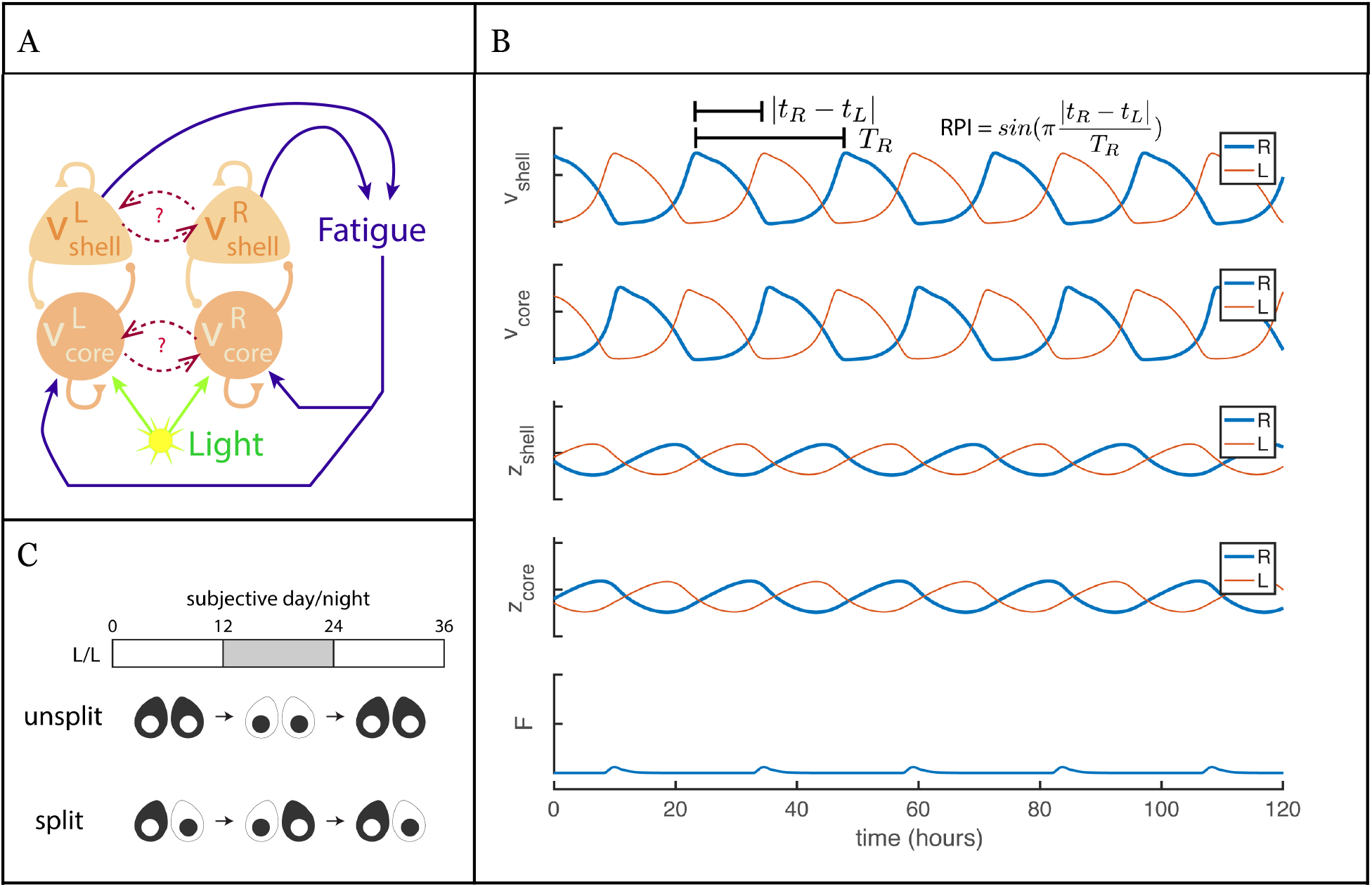
Bilateral SCN model produces splitting activity. A. Circuit schematic of bilateral SCN model showing the four activity nodes (*υ*^*R*^_*shell*_, *υ*^*L*^_*shell*_, *υ*^*R*^_*core*_, *and v*^*L*^_*core*_) and their connectivity. Fatigue is a readout of the *υ*^*R*^_*shell*_ and *υ*^*L*^_*shell*_ nodes. Light and Fatigue are inputs to the *υ*^*R*^_*core*_ and *υ*^*L*^_*core*_ nodes. B. Bilateral SCN model in split mode under constant dark conditions (L = 0). The two hemispheres are connected only by a shared Fatigue signal that enters the *υ*^*R*^_*core*_ and *υ*^*L*^_*core*_ nodes. Units on y-axes are arbitrary. The calculation of Relative Phase Index (RPI) is illustrated by annotations on the *υ*_*shell*_ traces (top panel). C. Schematic of split (bottom) and unsplit (middle) activity over the course of subjective day/night cycles (top). Adapted from (Butler et al., 2012).

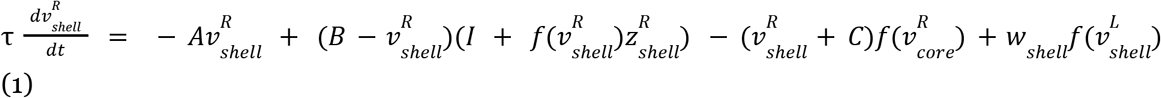

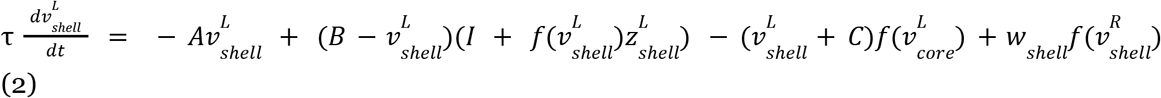

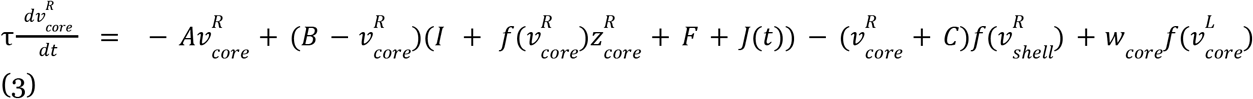

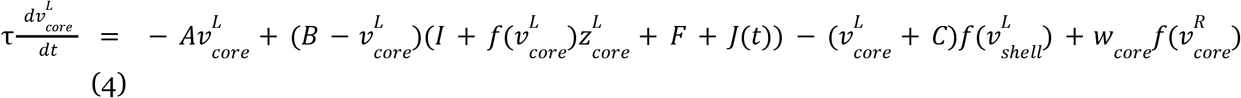

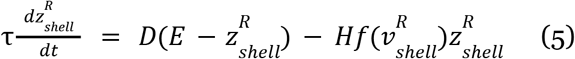

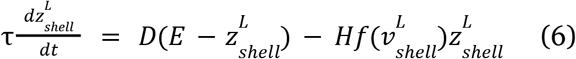

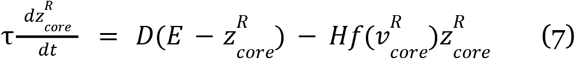

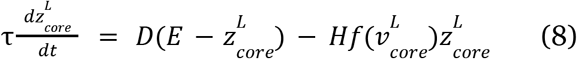

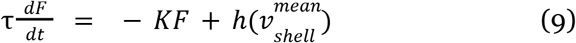

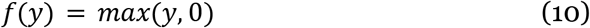

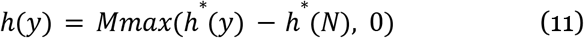

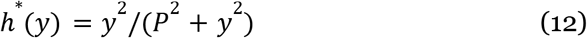

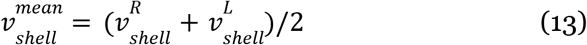

*J(t)* is the light input to the core nodes. It defaults to a constant value equal to light intensity, *L*, in constant light or constant dark conditions. In light/dark (LD) conditions, *J(t)* is a square wave signal that oscillates between 0 and *L* with a 50% duty cycle.

Our model has three distinct ways of connecting the left and right hemispheres: through the shared Fatigue signal, through shell to shell coupling (w_*shell*_), and through core to core coupling (w_*core*_). In the default version of our model, contralateral coupling across the shells or cores is “off” (i.e. w_shell_ = w_core_ = 0) and the two hemispheres are connected only via the shared Fatigue signal.

We retain most of the parameter values from Carpenter and Grossberg, 1984 (see Table). Our time constant, τ, was tuned to produce approximately 24 hour periods in constant dark (DD) for the *υ*_*core*_ and *υ*_*shell*_ nodes in the default bilateral model without explicit inter-hemisphere connections (i.e. w_shell_ = w_core_ = 0).

**Table 1.**
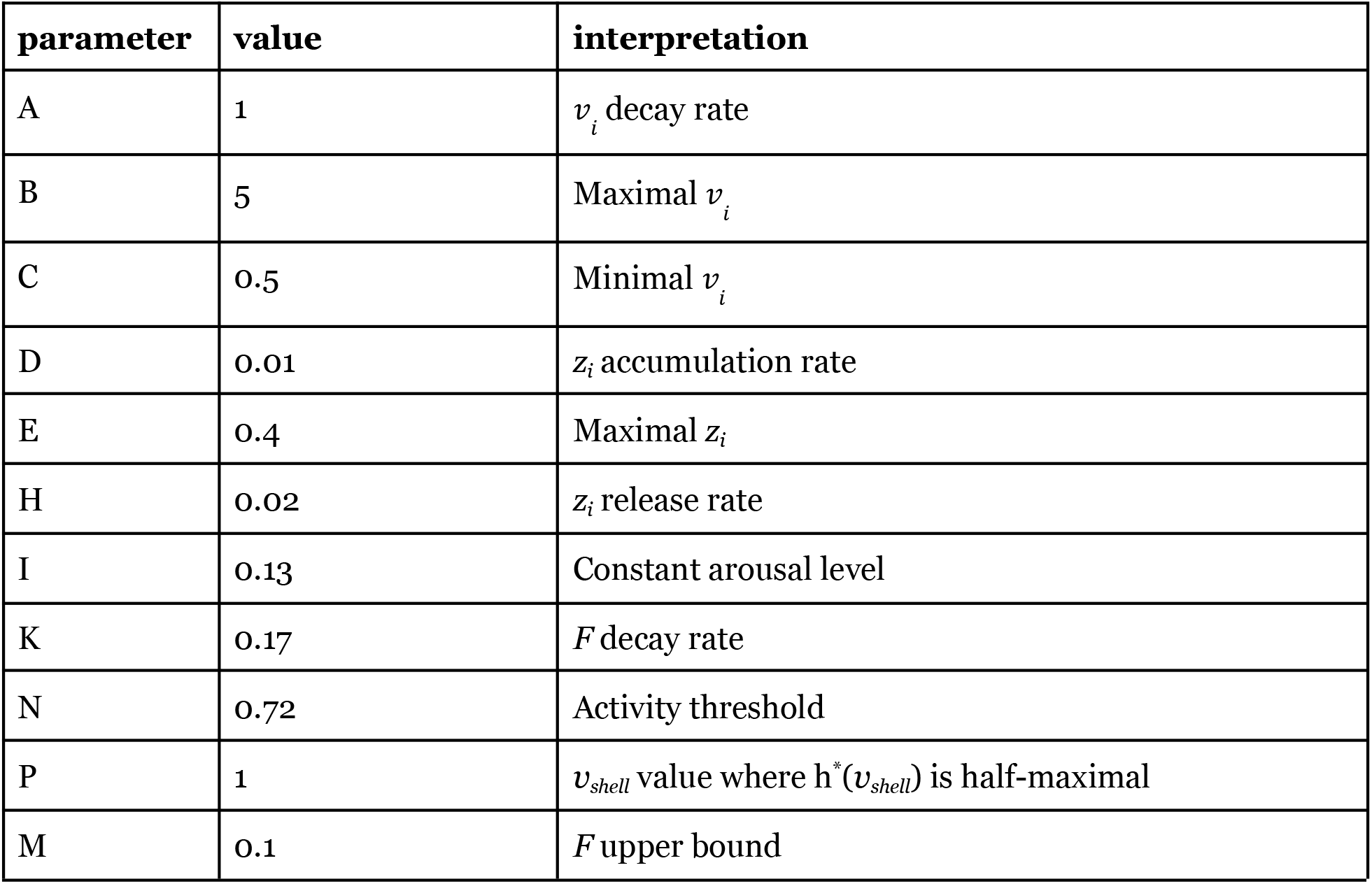
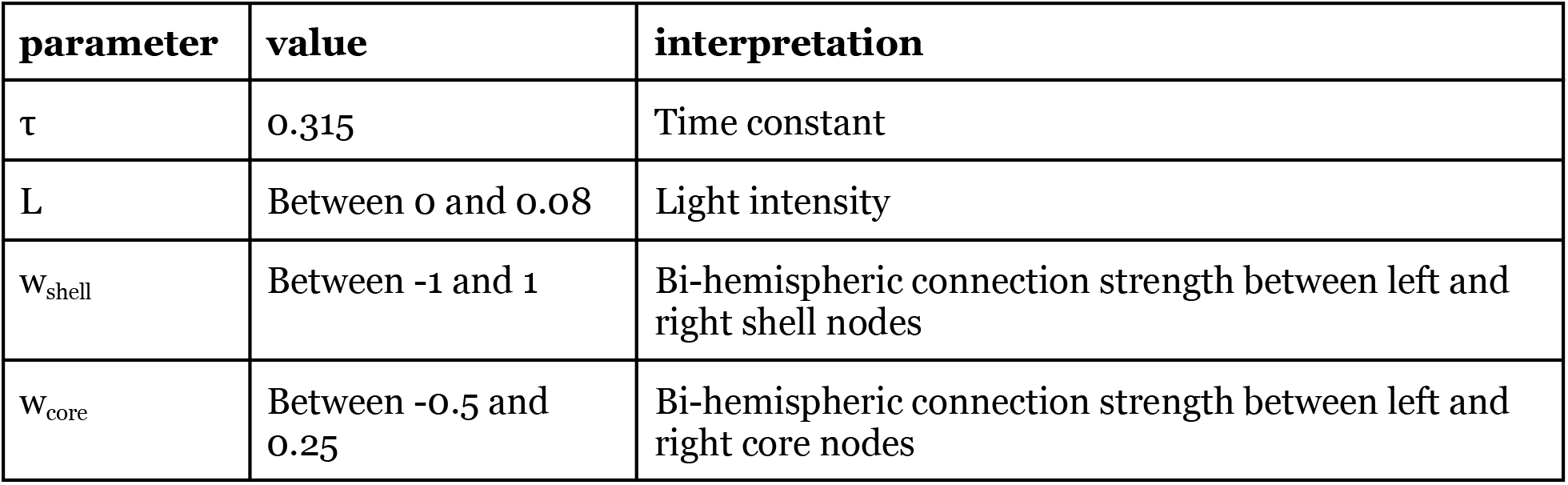
Model parameters.

| parameter | value | interpretation |
| --- | --- | --- |
| A | 1 | $v_i$ decay rate |
| B | 5 | Maximal $v_i$ |
| C | 0.5 | Minimal $v_i$ |
| D | 0.01 | $z_i$ accumulation rate |
| E | 0.4 | Maximal $z_i$ |
| H | 0.02 | $z_i$ release rate |
| I | 0.13 | Constant arousal level |
| K | 0.17 | $F$ decay rate |
| N | 0.72 | Activity threshold |
| P | 1 | $v_{shell}$ value where $h^*(v_{shell})$ is half-maximal |
| M | 0.1 | $F$ upper bound |
| $\tau$ | 0.315 | Time constant |
| L | Between 0 and 0.08 | Light intensity |
| $w_{\text{shell}}$ | Between -1 and 1 | Bi-hemispheric connection strength between left and right shell nodes |
| $w_{\text{core}}$ | Between -0.5 and 0.25 | Bi-hemispheric connection strength between left and right core nodes |

In the gated pacemaker model, the core and shell nodes oscillate at the same frequency with a shifted phase, otherwise there are no oscillations. The gating variables generally oscillate at the same frequency as the core and shell nodes when oscillations are present.

### Simulations

The model was implemented and analyzed using custom-written code in Matlab, Mathematica, and XPPAUT on a Mac Powerbook Pro or a Mac Studio. Simulations of model activity time series were performed in Matlab using the built-in ODE solver for non-stiff systems (ode45) based on Runge-Kutta integration of equations 1-9. Time is in units of hours. The maximum integration step size was bounded to 10 hours (MaxStep=10). The optimal tolerances depended on the simulation length and were set to RelTol=1e-5 and AbsTol=1e-7 for the 3000 hour simulations. For shorter simulations, RelTol=1e-4 and AbsTol=1e-6 was sufficient.

The first 500 hours of simulated time were discarded to allow the system to reach steady state. The model activity variables have arbitrary units. In all figures of model activity time series, scale bars indicate a consistent quantity across instances of the variable plotted. Zero is denoted by a small horizontal bar.

Simulated trials of model activity were distinguished by distinct initial conditions. Initial values for each of the 9 variables were independently drawn from uniform distributions with support as follows:

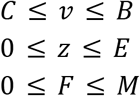

For each trial, the same initial condition was used for one sweep of the independent variable. Each combination of initial condition and independent variable value was simulated independently such that all combinations had the first 500 hours of simulated time discarded before being analyzed. The remaining 1200 hours of simulated time for each trial was analyzed (3000 hours of simulated time in the latency to switch trials). N = 100 trials, unless otherwise stated.

### Analyses of model activity

Oscillation periods were quantified using Fast Fourier Transform applied to the activity waveforms, *υ*, of the shell and/or core nodes. The periods among the four activity waveforms were typically identical regardless of whether the system was in the split or unsplit mode.

To quantify the relative phase between the right and left nodes, we utilized a Relative Phase Index, Φ, as illustrated in Fig. 1B where T_R_ is the period of signal R, and |t_R_ - t_L_| is the average absolute difference between the time of peaks of signals R and L:

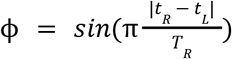

The Relative Phase Index varies between 0 and 1 such that two signals perfectly in phase will have a Relative Phase Index of 0 and two signals that are in perfect anti-phase will have a Relative Phase Index of 1.

A bifurcation analysis was performed on the bilateral system using XPPAUT.

## Results

### Split activity is dependent on Fatigue coupling and light conditions

Our model is an extension of the model by Carpenter and Grossberg (1984) with one half-center oscillator formed by two nodes. Here, we have two half-center oscillators, each one representing a hemisphere of the bilateral SCN (Fig. 1A). The behavior resulting from connecting two half-center oscillators, each with their own interconnected core and shell nodes, is not necessarily predictable and may be dependent on how they are connected. The first point of connection between the two hemispheres is the Fatigue signal, *F*. The Fatigue signal is a readout of the activity of the shell node and it is fed back into the core node. It represents an integration of the organism’s metabolic or locomotor activity (Carpenter and Grossberg, 1984). In our bilateral model, the Fatigue signal is the average activity of the left and right shell nodes (see eqn. 9) since having independent Fatigue signals for the right and left hemispheres of the SCN model would be difficult to justify given that the model represents the activity of one animal. The left and right core nodes thus receive identical feedback that is dependent on the combined shell activity.

When the left and right hemispheres share a Fatigue signal, the two hemispheres exhibit synchronized split activity (Fig. 1B,C). The right and left shell nodes are in antiphase with each other and the right and left core nodes are also in antiphase with each other. The right shell oscillates in phase with the left core, and the left shell oscillates in phase with the right core. The split activity emerges immediately when the right and left hemispheres are connected by a shared Fatigue signal under constant dark conditions (light intensity = 0), and it is robust to different initial conditions. We next asked what conditions can produce unsplit activity in the bilateral SCN model.

When we drive the model with 24-hour light/dark cycles where a 12-hour pulse of light is followed by a 12-hour period of darkness, we observe the unsplit mode where the left and right shell oscillate in phase, and the left and right core oscillate in phase, while the core and shell oscillate in antiphase within their respective hemisphere (Fig. 2A). These results depend on the intensity of the light pulses: higher intensity pulses are better able to entrain the model to the unsplit mode (Fig. 2B, bottom panel). Lower light intensities (L < 0.04) produce phase-shifted or split rhythms. The period of oscillations follow the 24-hour period of the light/dark cycles regardless of whether the model is split or unsplit in these lighting conditions (Fig. 2B, top panel).

**Figure 2.**
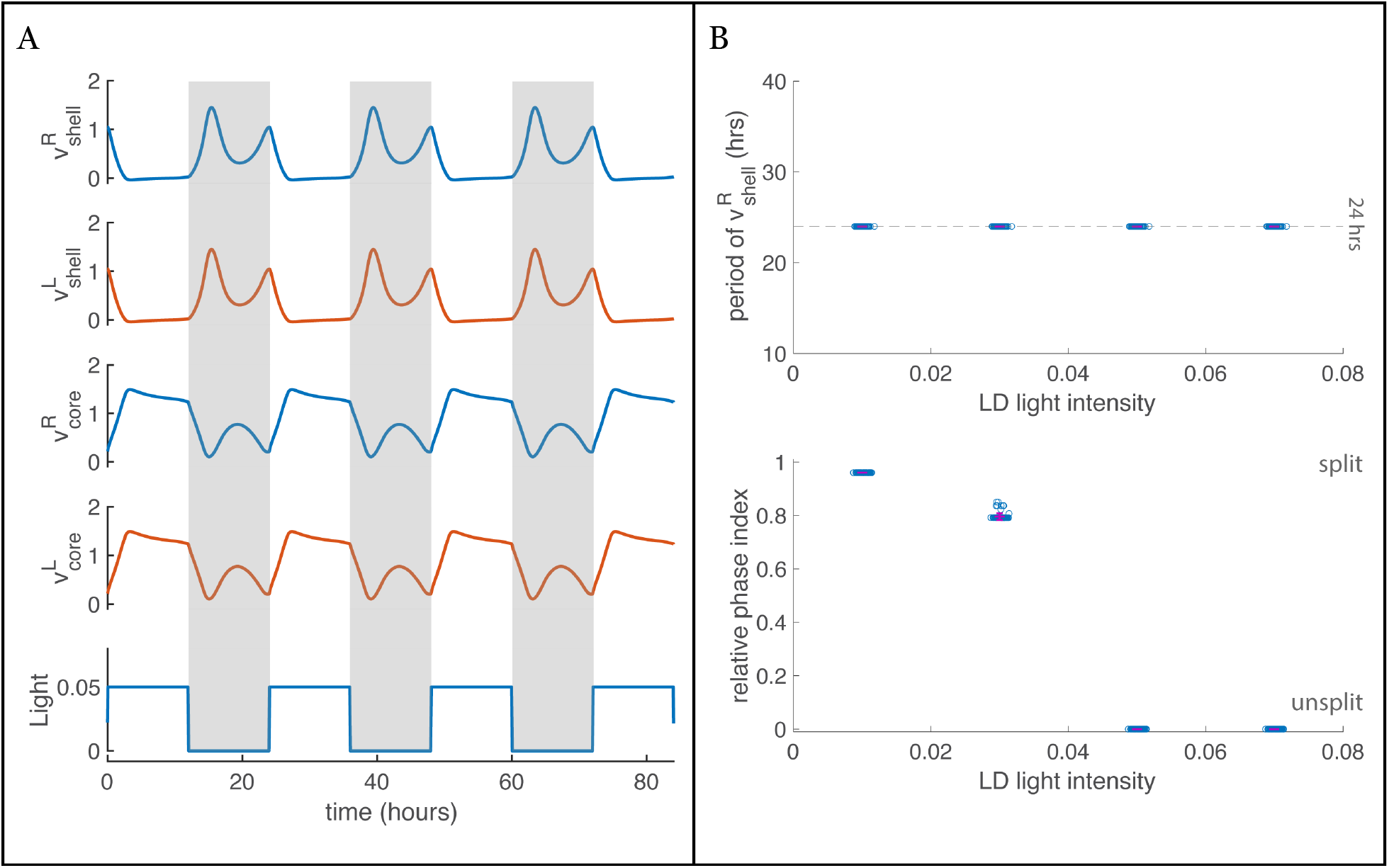
Model entrains to L/D cycles but only settles into unsplit activity for higher light intensities (>0.04). The two hemispheres are connected only by a shared Fatigue signal that enters the right and left core nodes (*υ*^*R*^_*core*_ and *υ*^*L*^_*core*_). A.The activity traces over time for the model entrained by light/dark cycles (light intensity, L =0.05). Gray highlights indicate dark periods. The hamster is a nocturnal animal and the shell nodes, *υ*^*R*^_*shell*_ and *υ*^*L*^_*shell*_, are most active during the night. B.Period of *υ*^*R*^_*shell*_ maintains the period of the 24-hour light/dark cycles (top). The relative phase between the *υ*^*R*^_*shell*_ and *υ*^*L*^_*shell*_ nodes is dependent on the light intensity of the light/dark cycles (bottom). Each point is a trial. There were 100 trials for each light intensity value, L (0.01, 0.03, 0.05, 0.07). Horizontal magenta bars denote the mean, vertical magenta bars denote the standard deviation.

At intermediate light intensities (L = 0.03), the relative phase is somewhat sensitive to the initial conditions (Fig. 2B, bottom panel) and there does not appear to be a stable split or unsplit regime. In contrast, the relative phase index is not sensitive to initial conditions when light intensity is low (L = 0.01), and the split mode is stable. When light intensity is high (L = 0.05, 0.07), the relative phase index is once again insensitive to initial conditions but the unsplit mode is stable. Therefore, light intensity may be a bifurcation variable whose adjustment could change the dynamics of the system.

In the classical splitting experiments, hamsters transitioned into split behavior after being kept in constant light conditions. We therefore next investigated the effect of driving the system by constant light of fixed intensity (Fig. 3). We found that the split mode was generally dominant across constant light conditions of different intensities. The exception to this was observed at light intensities in the middle range (0.03 through 0.06) where the relative phases were not consistent across trials – indicating sensitivity to initial conditions (Fig. 3 bottom). Indeed, some trials in the mid-intensity range resulted in unsplit activity with the period also decreasing to approximately 18-hours. Apart from this behavior in the mid light intensities, the oscillation period increased with increasing light intensities (Fig. 3 top).

**Figure 3.**
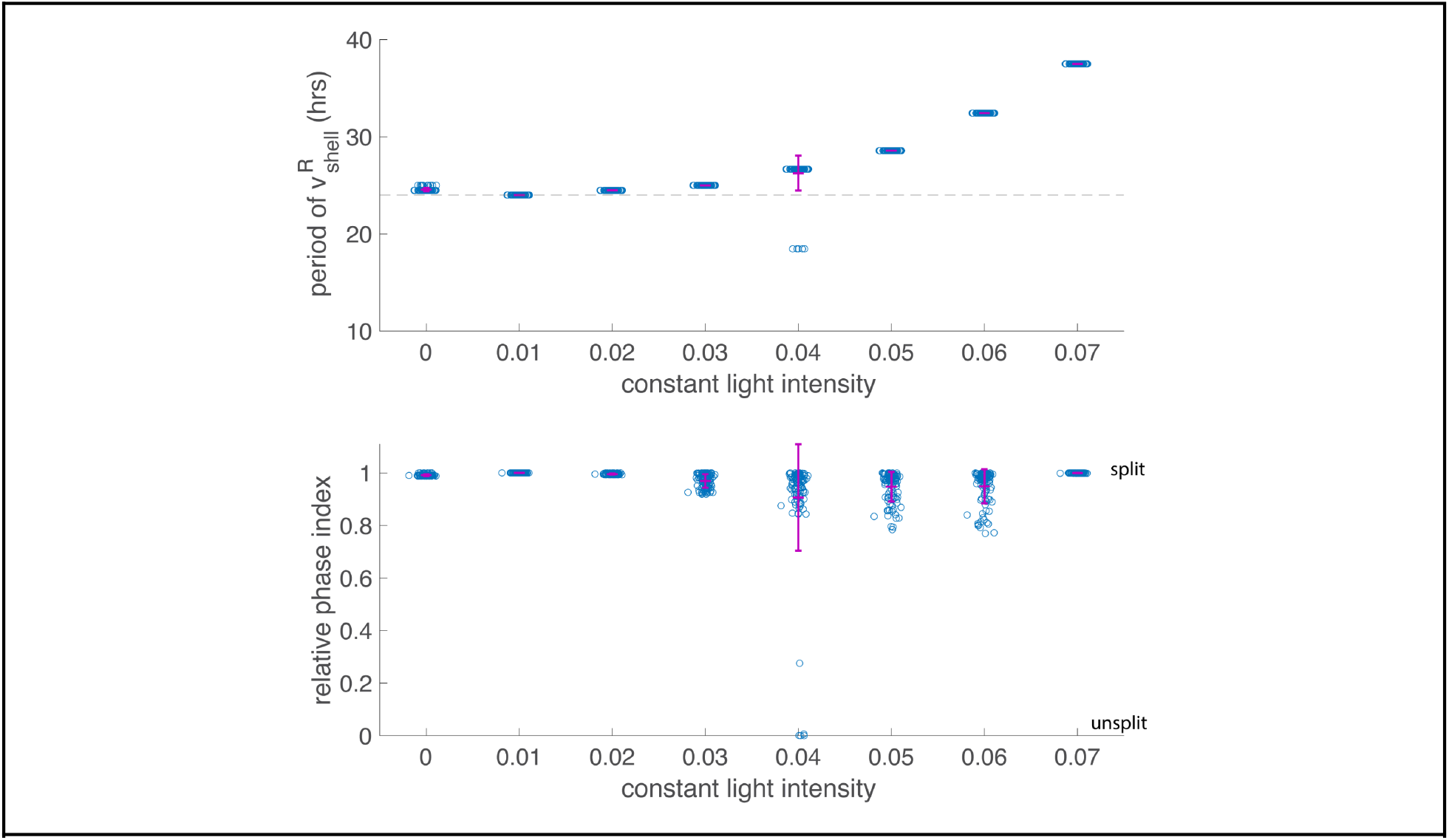
Period of *υ*^*R*^_*shell*_ (top) and relative phase between the *υ*^*R*^_*shell*_ and *υ*^*L*^_*shell*_ nodes (bottom) in simulations of the model in constant light conditions. The two hemispheres are connected only by a shared Fatigue signal that enters the *υ*^*R*^_*core*_ and *υ*^*L*^_*core*_ nodes. Note that we have D/D conditions when light intensity is zero. Each point is a trial. There were 100 trials for each light intensity value, L (0, 0.01, 0.02, 0.03, 0.04, 0.05, 0.06, 0.07). Horizontal magenta bars denote the mean, vertical magenta bars denote the standard deviation.

In hamster splitting experiments, the transition from unsplit to split behavior was not immediate, but emerged after some time in constant light, usually weeks. The intensity of the constant light affected the latency to split such that brighter constant light was more likely to lead to an earlier onset of splitting (Pickard et al., 1993). In our model, we observed a latency to split if the model was entrained to light/dark cycles before being switched to constant light (Fig. 4). Figure 4A shows the time course of the model nodes that are first entrained with several 24-hour cycles of light and dark pulses (light intensity, L = 0.05) and then exposed to total darkness (constant at an intensity of zero). As the model is transitioned to total darkness, the core and shell activity waveforms immediately change shape and decrease in amplitude. Despite these changes in waveform, the right and left nodes remain in phase for several oscillations (approximately from 300 to 650 hours). The model then spontaneously switches to split activity where the left and right nodes are out of phase with each other (after ∼650 hours). The latency to split is dependent on the constant light intensity that follows the L/D cycles (Fig. 4B). When constant light intensity is in the lower end of the range, the latency to split gradually increases as constant light intensity increases up until a light intensity of 0.04. In the higher end of the constant light intensity range, the latency to split gradually decreases as constant light intensity increases. Thus, the trend in the latency to split in the higher range of constant light intensities, but not the lower range, follows experimental observations.

**Figure 4.**
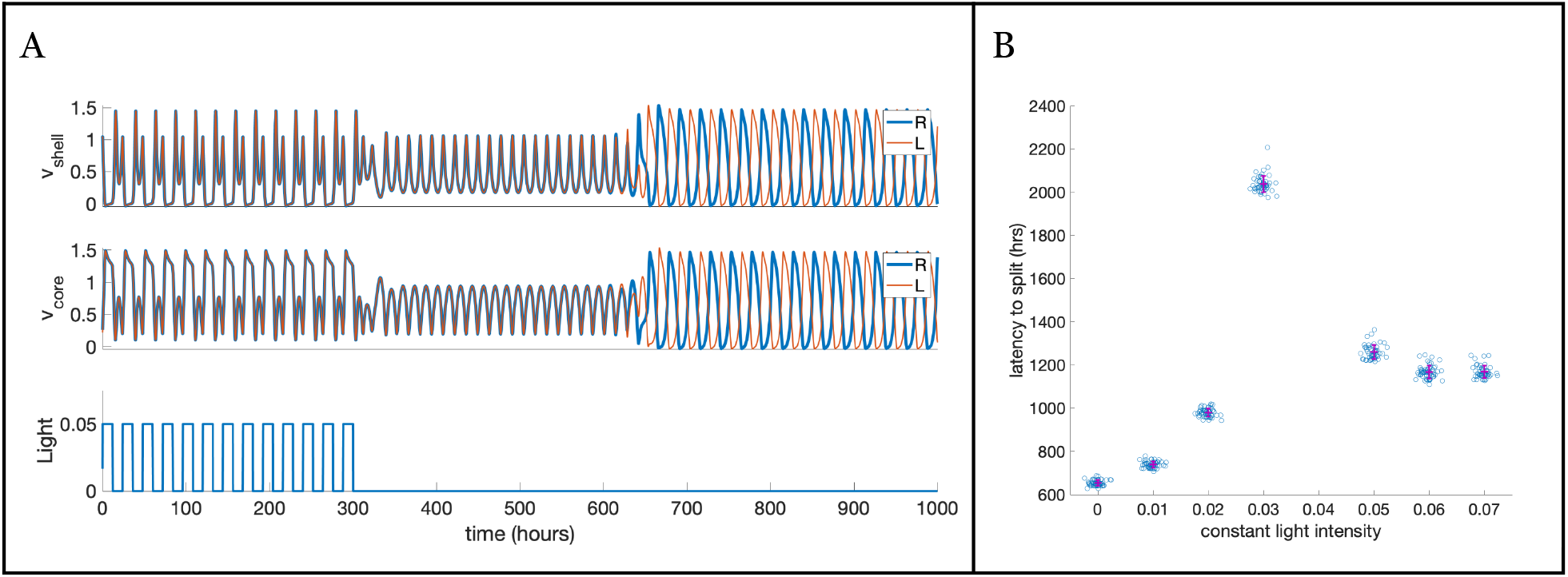
Latency to split. The two hemispheres are connected only by a shared Fatigue signal that enters the *υ*^*R*^_*core*_ and *υ*^*L*^_*core*_ nodes. A. Example trace showing model being entrained to L/D before constant light conditions. At 300 hours, the light stim switched to D/D and the model maintained unsplit activity for approximately 300 hours before spontaneously switching to split activity. B. The latency to split. Model entrained to L/D cycles with light intensity L = 0.05 from -500 to 300 hours as in (A). At 300 hours, the model was subjected to constant light. Latency to split first increases with increasing constant light intensity, then after L = 0.04 it begins to decrease with increasing constant light intensity

Our model displays split activity in constant darkness (Fig. 1B, 3, 4B) which is contrary to experimental findings. Even after entraining to light/dark cycles wherein the model displays unsplit activity, switching to constant darkness results in a relatively rapid switch to split activity (Fig. 4B). Biologically, the internal clock of the freerunning animal enables it to maintain approximately 24 hour synchronized activity rhythms even without light cues. The unilateral version of the model has freerunning 24 hour activity in constant darkness (see Carpenter and Grossberg, 1984). The hemispheres of our bilateral model maintain 24-hour activity cycles individually in constant darkness, but the right and left hemispheres are out of sync (see Fig. 3) which would cause the animal to have the 12-hour sleep/wake cycles that are characteristic of splitting. This inconsistency between our model and the experimental findings suggested that our model was missing an essential feature. Classic studies demonstrate the existence of commissural core-to-core and shell-to-shell fibers (van den Pol, 1980; Card et al., 1981; Leak et al., 1999; Moore et al., 2002), but little is known about the role that inter-hemispheric connectivity plays in the oscillatory dynamics that drive behavior. We therefore focus the remainder of our studies on investigating the effects of inter-hemispheric coupling in our model.

### Unsplit free-running activity requires excitatory contralateral connections

To determine whether and how inter-hemisphere connectivity affects the dynamics of the model, we swept across a range of coupling strengths for two configurations of bilateral coupling: mutually coupled core nodes, and mutually coupled shell nodes. In simulations with constant dark conditions, we found that unsplit activity required excitatory contralateral connections in either of the two configurations (Fig. 5). Positive coupling strengths tended to reduce the period of the oscillations relative to the oscillation periods observed with negative coupling strengths. However, larger magnitude core-to-core coupling strengths resulted in longer period oscillations up to a point. Beyond a shell coupling strength of 1.0 or a core coupling strength of 0.21, the model no longer produced oscillations. Thus, unsplit activity with a 24-hour period in constant darkness was only achieved with core node coupling at a strength of approximately 0.21, but unsplit activity with periods less than 24-hours were generally achieved with excitatory cross-coupling of either core or shell nodes. Our bilateral model predicts that excitatory inter-hemisphere connections are needed to produce unsplit activity in conditions of constant darkness.

**Figure 5.**
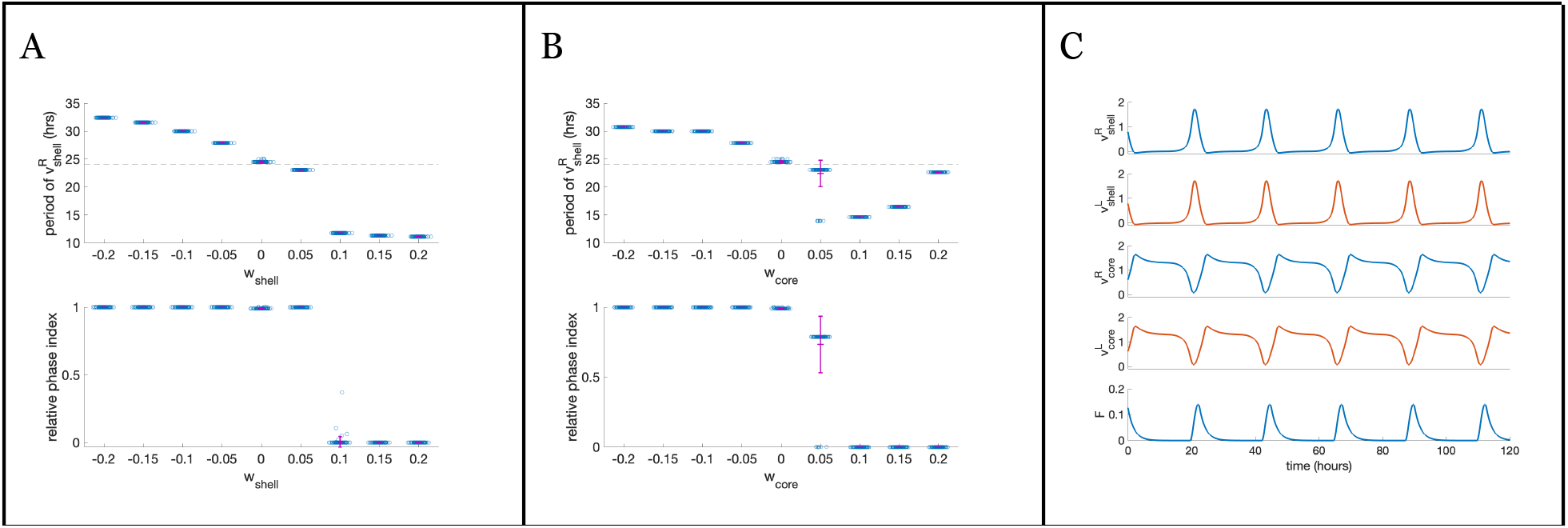
A. Period of *υ*^*R*^_*shell*_ (top) and relative phase between *υ*^*R*^_*shell*_ and *υ*^*L*^_*shell*_ (bottom) for sweeps through shell node contralateral coupling strengths, wshell, in constant dark conditions. B. Period of *υ*^*R*^_*shell*_ (top) and relative phase between *υ*^*R*^_*shell*_ and *υ*^*L*^_*shell*_ (bottom) for sweeps through core node contralateral coupling strengths, w_core_, in constant dark conditions. C. Example of model activity over time in constant dark (L = 0) with excitatory core node contralateral coupling strength, w_core_ = 0.2.

### Light intensity and cross-connectivity

Although excitatory contralateral connections enabled stable unsplit activity in constant dark conditions (Fig. 5), the model is sensitive to constant light when the core nodes are coupled. At a moderate core coupling strength (w_core_ = 0.2), the model ceased oscillations at light intensities greater than L = 0.01 (not shown). However, the model is able to entrain to light/dark cycles of moderate intensity when the core nodes are coupled with excitatory connections (not shown). Excitatory coupling between left and right shell nodes results in robust unsplit activity across the constant light intensity spectrum (Fig. 6A). Additionally, increasing light intensity has a tendency to increase the period of oscillations as it did for the model with no contralateral connections.

**Figure 6.**
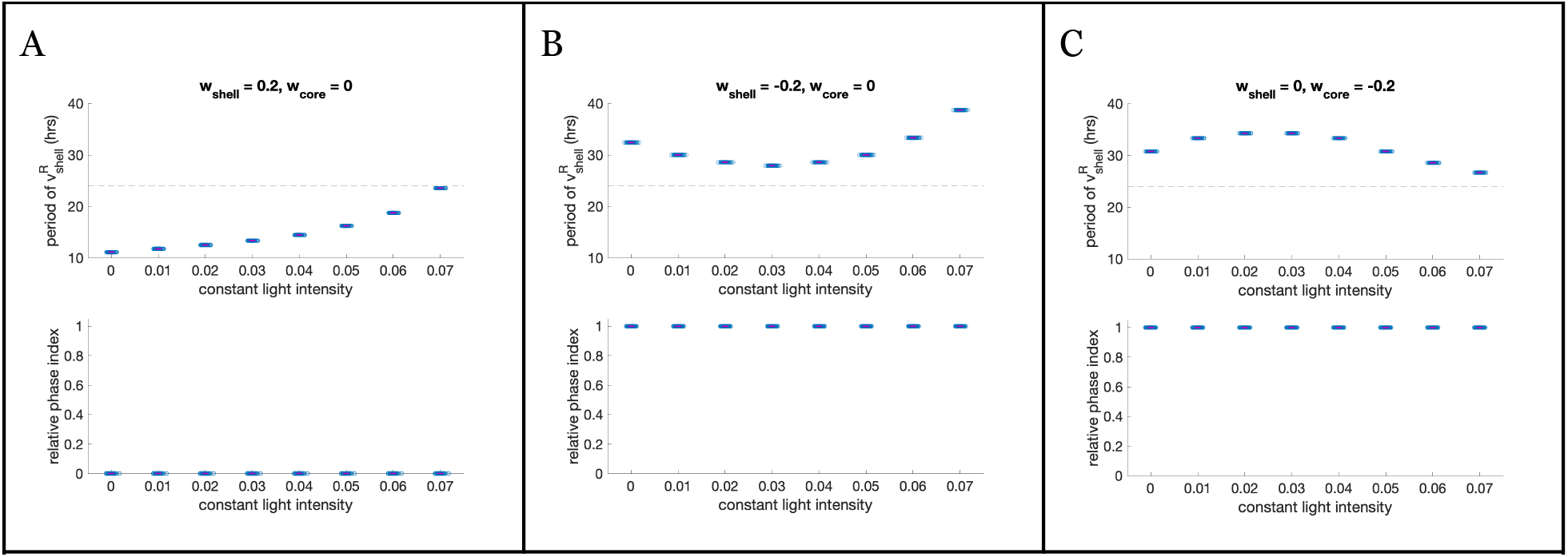
A. Period of *υ*^*R*^_*shell*_ (top) and relative phase between *υ*^*R*^_*shell*_ and *υ*^*L*^_*shell*_ (bottom) for sweeps through constant light intensity for model with excitatory inter-hemispheric shell coupling, *w*_*shell*_ = 0.2. B. Period of *υ*^*R*^_*shell*_ (top) and relative phase between *υ*^*R*^_*shell*_ and *υ*^*L*^_*shell*_ (bottom) for sweeps through constant light intensity for model with inhibitory inter-hemispheric shell coupling, *w*_*shell*_ = -0.2. C. Period of *υ*^*R*^_*shell*_ (top) and relative phase between *υ*^*R*^_*shell*_ and *υ*^*L*^_*shell*_ (bottom) for sweeps through constant light intensity for model with inhibitory inter-hemispheric core coupling *w*_*shell*_ = -0.2.

The unsplit activity displayed by the model with excitatory contralateral connections in constant dark conditions more closely resembles the activity one would observe in the biological system under constant dark conditions. Despite this, we did not observe a spontaneous switch to split activity in constant light, even when the model was entrained by light/dark cycles. This is because the excitatory connections stabilize the unsplit activity. Even a miniscule amount of excitatory cross-connectivity will push the latency to switch to later times until no switch can be observed within the time duration simulated. Given the lack of sensitivity to different initial conditions seen in Figure 6A, it is unlikely that the model with moderate to strong excitatory contralateral connections would exhibit such a switch, even for longer simulations.

Inhibitory contralateral connections, on the other hand, had the effect of stabilizing split activity rhythms. When shell nodes are coupled with inhibitory connections, split activity is observed for the full range of constant light intensities (Fig. 6B). The periods are generally longer than 24 hours and have a U-shaped relationship with the magnitude of constant light intensity. Likewise, when core nodes are coupled with inhibitory connections, the relative phase also shows split activity (Fig. 6C). The periods are also longer than 24 hours but with an upside-down U-shaped relationship with the magnitude of constant light intensity. In comparison to the model with no inter-hemispheric connections (Fig. 3), the split activity with inhibitory inter-hemispheric connections is far more robust to initial conditions and displays a more rigid anti-phase relationship between the right and left nodes within a trial.

### Bifurcation analysis

To better understand how the model dynamics depend on model parameters we next turn to bifurcation analysis. We focus this analysis on inter-hemisphere coupling strength and constant light intensity as our simulation results suggested that these are effective modes to transition model dynamics between split and unsplit modes.

First, we analyzed the effect of changing the connectivity strength coupling the left and right shell nodes (w_shell_). In the constant darkness condition, a one dimensional bifurcation analysis revealed that split and unsplit oscillations arise from two subcritical Hopf Bifurcations that face each other (Fig. 7A, points denoted by HB). The oscillations initially arising from the respective Hopf points are unstable (Fig. 7A, solid blue lines) but they become stable oscillations (Fig. 7A, solid green lines) through two Torus bifurcations occurring at w_shell_=0.0901 and w_shell_=0.0624. Since the two Hopf bifurcations face each other, the initial unstable oscillations arising from the respective Hopf bifurcations are enveloped by stable oscillations arising from the opposing bifurcation. This results in the model displaying stable split or unsplit oscillations for complementary w_shell_ values.

**Figure 7.**
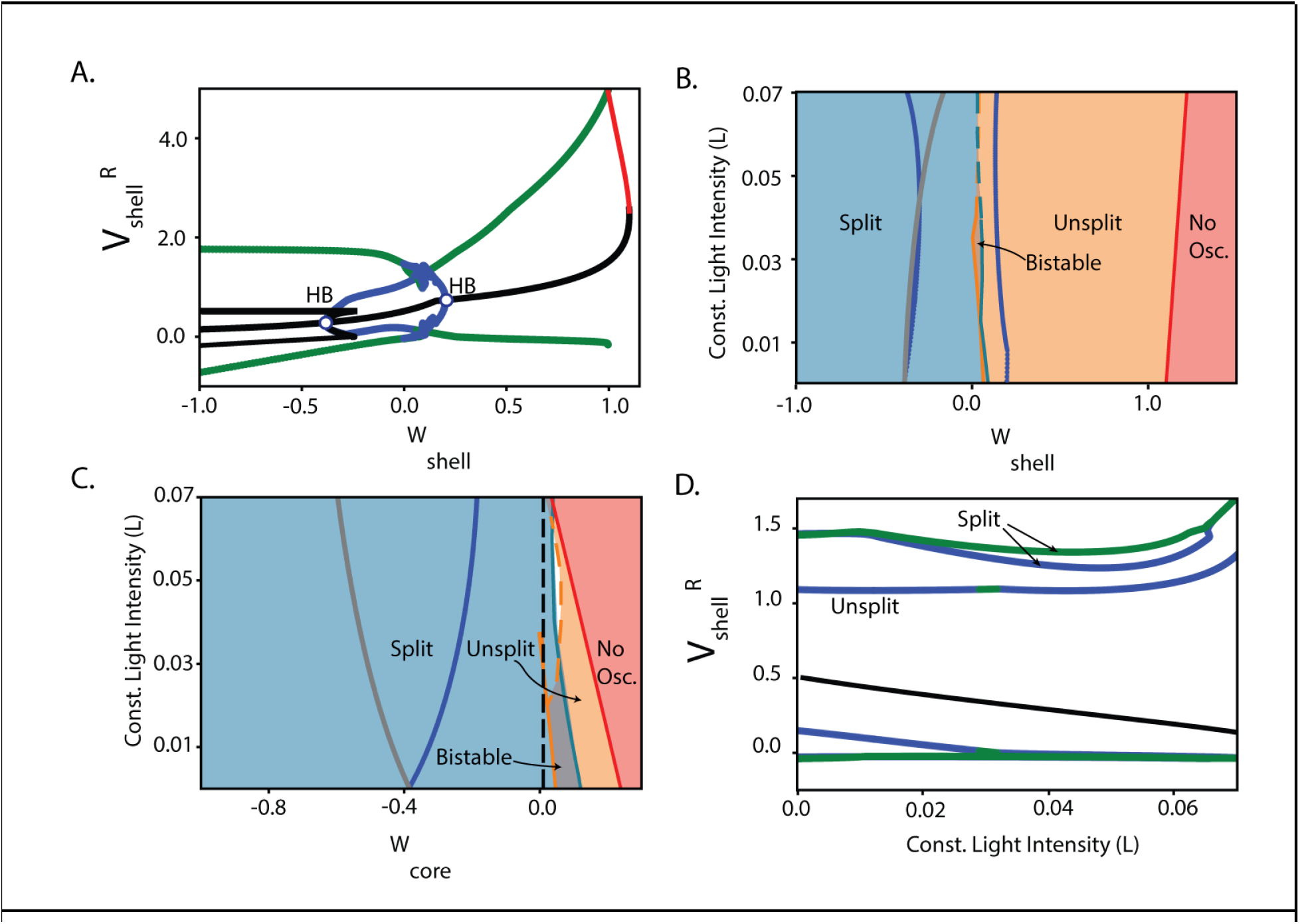
A.1D bifurcation for no light condition. Black and red lines indicate unstable and stable fixed points, respectively. Blue and Green lines indicate envelopes of unstable and stable oscillations, respectively. Hopf Bifurcation (HB) points are labeled with white dots. B.2D bifurcation of w_shell_ vs. constant light intensity. Blue lines indicate locations of Hopf bifurcations, grey line indicates bifurcation point where a single unstable fixed point branches into three unstable fixed points and red line indicates bifurcation point where unstable fixed point becomes stable. Regions of split, unsplit and no oscillations are indicated by blue, orange, and red shading respectively. Solid (dashed) border lines for unsplit and split regions indicate regions that are demarcated by the appearance of a torus (period doubling or other) bifurcation. C.2D bifurcation of w_core_ vs. constant light intensity. Colors are the same as in panel B. Dashed vertical black corresponds to panel D. The white region in the parameter space corresponds to a region that appears to display chaotic activity. D.1D bifurcation over constant light intensity at fixed w_core_ =0.01. Black solid line indicates an unstable fixed point. Blue and green lines indicate the envelope of split and unsplit stable oscillations. There are two types of split oscillations, one in which the *υ*^*R*^_*shell*_ and *υ*^*L*^_*shell*_ have the same amplitude and one in which they have different amplitudes.

The two torus bifurcations divided the space of model dynamics into split and unsplit regimes depending on the connectivity strength. It is also worth noting that there are two types of split oscillations, one in which the amplitude of the two hemispheres are the same and one in which they are different. For the following analysis, however, we will not distinguish between these two types of split modes. For connectivity weights such that w_shell_ <0.0901, we find the split mode where the period and amplitude of the oscillations of the two hemispheres remains relatively constant. For w_shell_>0.0624, we find the unsplit mode where the amplitude of the oscillations and period both increase as the w_shell_ parameter increases. Finally, for w_shell_ values close to 1, unsplit oscillations disappear by colliding with a stable fixed point (indicated by the red line in Fig. 7A) in a homoclinic bifurcation.

The two-dimensional bifurcation diagram in Figure 7B extends the previous analysis across a range of constant light intensities. For the range of constant light intensities (y-axis of Fig. 7B), the model generally behaves similarly with regions of split oscillations for negative and zero values of w_shell_, unsplit oscillations for positive values of w_shell_ and no oscillations as w_shell_ becomes large. There is also a small region of bi-stability with both unsplit and split oscillations appearing to be stable. For L>0.04, we find that the boundary of split and unsplit oscillations is no longer demarcated by a torus bifurcation but instead occurs at a Period Doubling bifurcation or other bifurcation (denoted by dashed teal/orange line in Fig. 7B).

Next, we look at the two dimensional bifurcation analysis for w_core_ vs. constant light intensity (Fig 7C). We again see that the unsplit mode arises through a subcritical Hopf bifurcation (blue line), the oscillations of which become stable through a torus bifurcation for low to medium levels of constant light intensity (solid orange line), or a period doubling bifurcation (dashed orange line) for medium to high levels of constant light intensity. These stable unsplit oscillations persist until the oscillations collide with the branch of stable fixed points (red line). Stable oscillations for the split regime occur for a large range of w_core_ values and are bounded below by the torus bifurcation (Fig. 7C, teal line). There are two small regimes of bi-stability (Fig. 7C, one is to the left of the teal line and to the right of the orange line, indicated by joint shading in both orange and teal, and the second is in the region where the top of the solid orange line protrudes into the teal region) where both unsplit and split modes coexist. However, these regimes disappear as constant light intensity is increased (Fig. 7C, moving up the y-axis).

Finally, we investigated what happens for fixed w_core_ values as the level of constant light intensity is varied. As an example, we consider setting w_core_=0.01 and look at the corresponding one dimensional bifurcation over the range of constant light intensities. This analysis (shown in Figure 7D, corresponding to the vertical dashed black line through Fig. 7C) reveals that the split mode is stable for all light intensity values, as indicated by the presence of the green line for the entire range of constant light intensities at this core connectivity strength. The unsplit mode, however, is mostly unstable except for a small region (0.029<=L<=0.032). This region corresponds to the small region of bi-stability that was discovered in Figure 7C. Together, panels 7B and 7C show that it is generally not possible to cross from a regime of stable split to unsplit modes by only changing the level of light in this model. The only exception to this would be to initialize the model on the split solution in a region of multi-stability and then move to the unsplit regime by increasing the magnitude of light intensity but this would require fine tuning in both the initial conditions as well as either of the inter-hemisphere coupling strengths.

## Discussion

Our study introduces a bi-hemispheric mathematical model of the SCN and demonstrates how the dynamics of the SCN may depend on factors such as external light conditions and hemisphere synaptic coupling strengths. Through simulation and bifurcation analysis, we provide a mechanistic interpretation of previously observed experimental phenomena and suggest new experiments that could further illuminate the circuit underlying the SCN.

We first used simulations to investigate the conditions under which the bilateral model produces split versus unsplit activity. We found that the bilateral model automatically has split activity unless there are excitatory inter-hemispheric connections or strong light/dark input. This finding suggests that excitatory contralateral connections are needed to explain the experimentally observed synchronization seen in freerunning behavior (see Fig. 5). We also found that adjusting the strength and polarity of the inter-hemispheric connections was more effective at transitioning the model from unsplit to split than adjusting the constant light intensity (compare Figs. 5 and 6), suggesting that plasticity of commissural connections may be involved in the splitting phenomenon.

To systematically evaluate the effect of inter-hemispheric connectivity and light input parameters, we next turned to bifurcation analysis. The analysis identified three regimes in the model corresponding to unsplit oscillations, split oscillations or the absence of oscillatory behavior altogether. While these three regimes existed when we varied coupling strength for either shell or core nodes, the allowable ranges of coupling strength and constant light intensity of these regimes varied significantly. For instance, our analysis revealed a larger range of coupling strengths for the shell nodes compared to the core nodes that results in unsplit activity, meaning that the unsplit activity is more robust to changes in synaptic strength. On the other hand, there is a more robust range of coupling strength of the core nodes relative to constant light intensity that supports a regime of bistability of unsplit and split modes. Furthermore, we found that there is no shell or core coupling strength, of those considered, for which the system changes from a region of purely unsplit to purely split regimes as constant light intensity increases. The bistability region identified in Fig. 7C does suggest that it is possible to account for this observation by setting appropriate initial conditions to produce the unsplit mode in total darkness and then transition to the split mode as constant light intensity increases. This, however, would require fine-tuning of the core coupling parameter as well as fine tuning in the initial condition. The proposed region of bistability would also suggest that it should be possible to switch between the unsplit and split regions in complete darkness through a change in initial conditions such as the injection of current into the core or shell nodes. While we only explore the inclusion of either shell or core coupling in the present analysis, it is possible that including both types of coupling may provide a more robust way to explain light-dependent transitions between split to unsplit behavior.

Our model is an expansion of a unilateral model of the SCN that explained the transition into the split rhythm as a frequency doubling of the core and shell cycles (Carpenter and Grossberg, 1985). While the classic splitting behavior does present as a doubling of the rest/wake cycle frequency (Pittendrigh, 1960b; National Research Council, 1971), the underlying activity of the SCN during splitting was not known at the time that Carpenter and Grossberg developed their model. Subsequent experiments revealed that the ipsilateral core and shell are in antiphase while the left and right hemispheres are also in antiphase, all while each of the four nodes oscillate with an approximate 24-hour period (de la Iglesia et al., 2000; Yan et al., 2005; Butler et al., 2012). This results in a doubling of rest/wake cycles as the animal is driven into a “split” brain state by the divergent activity of the left and right SCN (de la Iglesia et al., 2000). In a model with a finite number of phase oscillators, where half the population has the opposite phase as the other half, negative coupling terms were required for stable activity (Indic et al., 2007). Such a model captures either the phase-splitting of the left and right SCN or the phase-splitting of the core and shell, but it does not capture both simultaneously. Our model more accurately captures the dynamics of the split SCN than previous models that simulate the transition to the split activity as a frequency-doubling of any part of the SCN.

A model of two coupled oscillators formed by Wilson-Cowan-style populations of mutually coupled excitatory and inhibitory nodes exhibits the 4-node split (Kawato and Suzuki, 1980). As their “environmental” parameter is varied, the model transitions between split and unsplit activity similar to how our model transitioned between split and unsplit with changes in the inter-hemisphere coupling. Like the dual-oscillator Wilson-Cowan model, our bifurcation analysis also revealed that the split or unsplit oscillations stabilize via a torus bifurcation.

Similar bifurcations have been observed in other systems of weakly coupled oscillators (Borisyuk et al., 1995). Our model most closely resembles the excitatory to excitatory coupling between Wilson-Cowan oscillators described in Borisyuk et al. 1995 in that for very small positive coupling strength, the system displays a stable split mode and then transitions to a stable unsplit mode through a series of torus bifurcations that interact with each other. Unlike the Borisyuk et al. (1995) model which features interaction between explicit excitatory and inhibitory populations, our system has shell and core nodes that self-excite but inhibit each other as well as an additional Fatigue signal that affects the dynamics. As a result, the specifics of the interactions of the bifurcations that transition the system from split to unsplit modes differ.

In conclusion, we sought to model the 4-node split to investigate how light intensity and coupling strength affect the transition from unsplit to split activity. Our results suggest that excitatory inter-hemisphere SCN coupling may be responsible for stable, in-phase freerunning activity across the two hemispheres, and that plasticity or modulation of the inter-hemisphere connections can elicit 4-node split activity. Our reduced model does not capture the emergent dynamics that result from the thousands of individual oscillators that compose the SCN, instead lumping entire populations of the SCN into four distinct nodes. Future studies may test the role of inter-hemispheric connections in a more detailed model.

## Acknowledgements

We would like to thank Rae Silver, Duncan Foley, and Nico Foley for their early involvement in this study. This work was also supported in part by the Barnard-Columbia Summer Research Institute (SRI) and the Science Pathways Scholar Program (SP2) at Barnard College.

## Author Contributions

Contributions according to [CRediT criteria](https://www.sagepub.com/journals/information-for-authors/submitting-your-manuscript/credit).

| # | Role | Definition | author(s) |
| --- | --- | --- | --- |
| 1 | Conceptualization | Ideas; formulation or evolution of overarching research goals and aims. | GJG |
| 2 | Data curation | Management activities to annotate (produce metadata), scrub data and maintain research data (including software code, where it is necessary for interpreting the data itself) for initial use or later reuse. | GJG |
| 3 | Formal analysis | Application of statistical, mathematical, computational, or other formal techniques to analyze or synthesize study data. | GJG, KZ, JM, AL, JN |
| 4 | Funding acquisition | Acquisition of the financial support for the project leading to this publication. | GJG |
| 5 | Investigation | Conducting a research and investigation process, specifically performing the experiments, or data/evidence collection. | GJG, KZ, JM, AL, JN |
| 6 | Methodology | Development or design of methodology; creation of models. | GJG, KZ |
| 7 | Project administration | Management and coordination responsibility for the research activity planning and execution. | GJG |
| 8 | Resources | Provision of study materials, reagents, materials, patients, laboratory samples, animals, instrumentation, computing resources or other analysis tools. | GJG |
| 9 | Software | Programming software development; designing computer programs; implementation of the computer code and supporting algorithms; testing of existing code components. | GJG, KZ, JM, AL, JN |
| 10 | Supervision | Oversight and leadership responsibility for the research activity planning and execution including mentorship external to the core team. | GJG |
| 11 | Validation | Verification whether as a part of the activity or separate, of overall replication/reproducibility results/experiments/other research outputs. | GJG |
| 12 | Visualization | Preparation creation presentation published work specifically visualization/data presentation. | GJG, KZ |
| 13 | Writing – original draft | Preparation creation presentation published work specifically writing initial draft (including substantive translation). | GJG, KZ |
| 14 | Writing – review & editing | Preparation creation presentation published work by those from original research group specifically critical review commentary revision including pre-post-publication stages. | GJG, KZ, JM, AL, JN |

## Statements and Declarations

### Ethical Considerations

No human or animal subjects were included in this study.

### Consent to Participate

Not applicable

### Consent for Publication

N/A

### Data Availability

All data are computer-simulated and can be regenerated using the simulation code available in the SCN-split-model Github repository at https://github.com/Gutierrez-lab/bilateral-SCN-gated-pacemaker.

